# McAN: an ultrafast haplotype network construction algorithm

**DOI:** 10.1101/2022.07.23.501111

**Authors:** Lun Li, Bo Xu, Dongmei Tian, Cuiping Li, Na Li, Anke Wang, Junwei Zhu, Yongbiao Xue, Zhang Zhang, Yiming Bao, Wenming Zhao, Shuhui Song

**Author notes:** To whom correspondence should be addressed. (Song S), (Zhao W), (Bao Y), (Zhang Z), (Xue Y). Equally contributed (Lun Li, Bo Xu, Dongmei Tian). **Supplementary information:** Supplementary material and figures are available at *Bioinformatics* online.

## Abstract

**Summary:** Haplotype network is becoming popular due to its increasing use in analyzing genealogical relationships of closely related genomes. We newly proposed McAN, a minimum-cost arborescence based haplotype network construction algorithm, by considering mutation spectrum history (mutations in ancestry haplotype should be contained in descendant haplotype), node size (corresponding to sample count for a given node) and sampling time. McAN is two orders of magnitude faster than the state-of-the-art algorithms, making it suitable for analyzation of massive sequences.

**Availability:** Source code is written in C/C++ and available at https://github.com/Theory-Lun/McAN and https://ngdc.cncb.ac.cn/biocode/tools/BT007301 under the MIT license. The online web service of McAN is available at https://ngdc.cncb.ac.cn/ncov/online/tool/haplotype. SARS-CoV-2 dataset are available at https://ngdc.cncb.ac.cn/ncov/.

## 1 Introduction

Haplotype network is fundamental to determine and visualize genealogical relationships of population genomes, thereby playing important roles in tracing the evolution and migration of diverse species (Bandelt, et al., 1995; Yue, et al., 2018). Typically, haplotype network construction algorithms infer ancestry-descendance relationships among haplotypes according to the distance matrix between haplotypes. Over the past decades, several algorithms have been proposed for haplotype network construction, such as minimum spanning network (MSN) (Bandelt, et al., 1999), median-joining network (MJN) (Bandelt, et al., 1999) and Templeton Crandall and Sing algorithm (TCS) (Templeton, et al., 1992), which have been applied to a variety of genome sequences, including virus, human mitochondria (Bandelt, et al., 1995), plant chloroplast (Yue, et al., 2018) and mammalian Y chromosome (Felkel, et al., 2019). Haplotype network is extensively used in virus studies, with the aim to identify possible transmission routes of virus particularly during the COVID-19 pandemic (Kemenesi, et al., 2020; Sekizuka, et al., 2020; Song, et al., 2020; Zhao, et al., 2020). Unfortunately, the existing algorithms have significant limitations as follows. First, they ignore several important features when constructing haplotype network, such as sampling time, number of sequences in haplotypes, and mutation spectrum history, which can mislead the tracing of evolution/transmission routes. Second, they generate undirected networks, accordingly unable to reflect the evolutionary directions. Third, all of them are time-consuming and memory-intensive, incapable to deal with massive datasets, like millions of SARS-CoV-2 sequences to date. The time complexities of MSN, MJN and TCS are *O*(*m*^2^), *O*(*m*^2.2^) and *O*(*m*^5^), respectively, where *m* is the number of haplotypes. In practice, MJN takes more than three days to construct haplotype networks for 1,000 SARS-CoV-2 sequences, as it calculates and stores distance matrix between any two haplotypes. To address these issues, here we present McAN (Minimum-cost Arborescence Network), a new haplotype network construction algorithm that is capable to build a directed, rooted tree spanning all vertices with minimum cost and considers genome-wide mutation spectrum features and epidemiological characteristics. As testified on large-scale SARS-CoV-2 genome sequences, McAN achieves accurate and efficient haplotype network construction.

## 2 Methods

McAN is built based on minimum-cost arborescence for haplotype network construction, a well-known graph theory widely used in multiple applications, e.g., irrigation system (Dutta and Mishra, 2012), multiple-people tracking in video sequences (Henschel, et al., 2015). In McAN, haplotype network is represented by a directed, rooted arborescence, in which each haplotype is a cluster of sequences with same mutations, each edge reflects directed ancestor-descendant relationship between haplotypes, and optimal arborescence is determined by minimum distances of all summed edges. Specifically, when constructing haplotype network, McAN takes good account of mutation spectrum history and assumes that mutations in descendant haplotype should contain in its ancestry and larger haplotype corresponding to more sequences is more likely to be an intermediate ancestor node (Figure S1). If sampling time is available, ancestral mutants are believed to be earlier sampled than derived mutants. All haplotypes are sorted by mutation count and sequence count in descending order and the earliest sampling time (if available) in ascending order (Figure S2A). Based on this sorting, the closest ancestor is determined and minimized for each haplotype (Figure S2B). Moreover, to save memory and running time, McAN calculates distances between adjacent haplotypes instead of any two haplotypes (Figure S2C). More details about McAN are provided in Supplementary materials (Text S1).

## 3 Results

We evaluate the accuracy of McAN by comparison with existing popular algorithms MSN, MJN, and TCS. On a simulated dataset with 1,023 sequences (see Text S2), we find that the AUC (area under the curve) of McAN, MSN, MJN, and TCS are 0.998134, 0.999972, 0.996214, and 0.992469, respectively, showing that McAN, albeit slightly lower than MSN, outperforms the other two algorithms. Moreover, we test on a real dataset of 482 SARS-CoV-2 genome sequences isolated from the *Diamond Princess* cruise as well as multiple countries/regions globally as detailed in (Sekizuka, et al., 2020). Strikingly, unlike the other three algorithms, McAN produces a direct evolutionary route from the reference (MN908947.3) to the *Diamond Princess* cluster (Figure S3), agreeing well with a finding that SARS-CoV-2 dissemination on the *Diamond Princess* cruise is originated from a single introduction event (Sekizuka, et al., 2020). Furthermore, we tested the accuracy of McAN for tracing the evolution of SARS-CoV-2 during the development of the global pandemic of COVID-19. The haplotype network constructed by McAN using 130 major haplotypes of sublineages of L and S lineages from 121,618 SARS-CoV-2 genomes (Tang, et al., 2021) shows a distinct delineation among these sublineages (Figure S4), suggesting that McAN can track the evolution of SARS-CoV-2 precisely. Except for SARS-CoV-2, McAN also construct reasonable haplotype networks for other kinds of viruses including Monkeypox virus (MPXV) (Gigante, et al., 2022) and human influenza A viruses (IAV) (Patrono, et al., 2022) (Figure S5).

We test the running performance of McAN using 1,124,837 SARS-CoV-2 genome sequences retrieved from RCoV19 (Gong, et al., 2020; Song, et al., 2020; Zhao, et al., 2020) as of 26 July 2021 on a personal laptop (Intel Core i5-6200U CPU and 8GB memory running with Ubuntu 20.04 operating system). We first assemble a small dataset by randomly sampling SARS-CoV-2 sequences. When the number of sequences ranges from 200 to 1,000, McAN consistently obtains higher efficiency, running about 1,000 times faster than TCS and MJN and 100 times faster than MSN (Figure 1A). Moreover, we test on a larger dataset with different sequence counts ranging from 100 thousand to one million (which shows a significant positive correlation with haplotype count; Figure S6). Noticeably, even when sequence count reaches 1 million, McAN just takes less than 20 minutes (Figure 1B), together demonstrating that McAN is well fitted to construct haplotype network with largescale dataset.

**Fig. 1.**
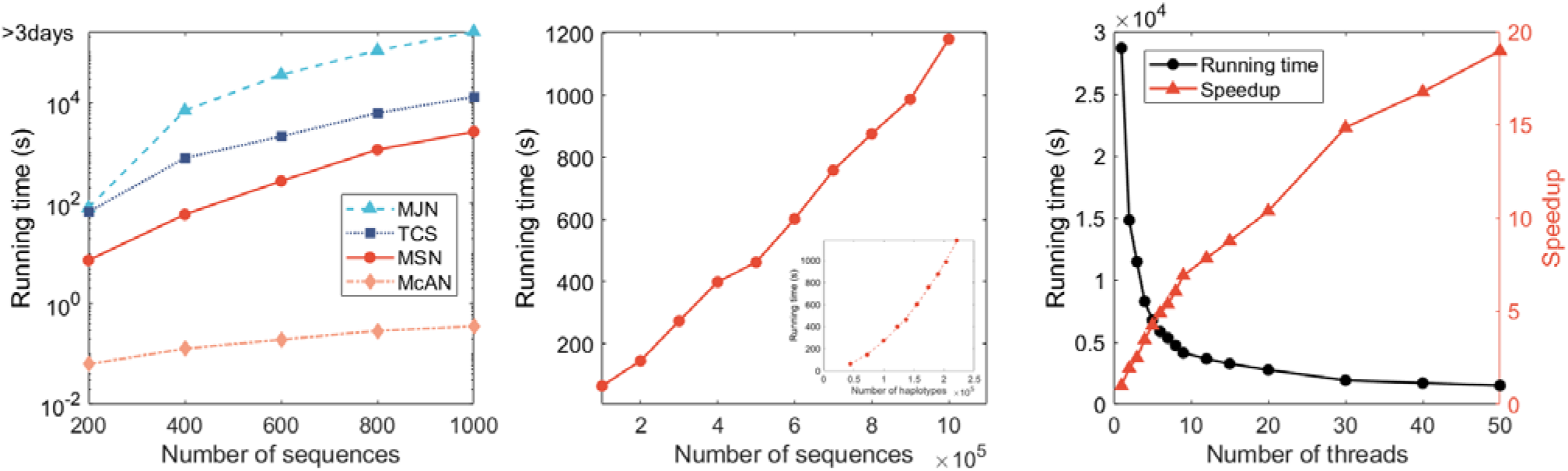
Performance of McAN by comparison with existing algorithms. (A) Comparison among MSN, MJN, TCS, and McAN in terms of running time wit**h** the number of sequences ranging from 200 to 1,000 (single thread); (B) Running time of McAN with the number of sequences ranging from 100,000 to 1,000,**000** (single thread); (C) Running time of McAN with number of threads ranging from 1 to 50 for 4,990,399 sequences.

Facing to millions of SARS-CoV-2 sequences, we further develop a parallel version of McAN. We test the performance of paralleled McAN on a server (2 Hygon C86 7185 32-core Processor, 512GB memory with CentOS Linux release 7.4.1708 (Core) operating system) using 4,990,399 SARS-CoV-2 sequences from RCoV19 as of 20 April, 2022. When the number of threads ranges from 1 to 50, the running time decline rapidly from 28,696.5s to 1,511.9s (Figure 1C).

## 4 Conclusion

McAN is an ultrafast new haplotype network construction algorithm, enabling real-time molecular tracing of pathogens for pandemics (e.g., COVID-19). As testified on multiple datasets, McAN outperforms existing algorithms and achieves higher accuracy and efficiency in haplotype network construction, suitable for massive data analysis in the big data era.

## Acknowledgements

The authors thank the members of China National Center for Bioinformation for providing SARS-CoV-2 mutation data and metadata.

## Funding

This research was funded by grants from the National Key Research & Development Program of China (2021YFF0703703, 2021YFC0863300 to S.H.S.), the Strategic Priority Research Program of the Chinese Academy of Sciences (XDB38060100 to Y.M.B.), the National Natural Science Foundation of China (32170678 to W.M.Z.), Youth Innovation Promotion Association of CAS (2017141 to S.H.S.) and the Beijing Nova Program (Z211100002121006 to L.L.).

### Conflict of Interest

none declared.

## Notes

### Competing Interest Statement

The authors have declared no competing interest.

