## Supplementary Material for "McAN: an ultrafast haplotype network construction algorithm"

**Text. S1. Haplotype network construction algorithm.**

The sequence accession ID and mutation pair were stored in a hash map, and sequences with the same mutations were clustered into a group, also called a haplotype. We inferred the direct ancestor-descendant relationships or immediate ancestor-descendant relationships for all haplotypes, which are represented by an arborescence (Gordon and McMahon, 1989) (called directed rooted tree and also called haplotype network in the field of bioinformatics). Specially, each haplotype is represented by a node or a vertex and each direct ancestor–descendant relationship is represented by a directed edge in haplotype network. And the reference sequence (users can customize any sequence) is designated as the root node of the haplotype network.

McAN factors three features into constructing haplotype network, including mutation spectrum history (mutations in ancestry haplotype should be contained in descendant haplotype), node size (corresponding to sample count for a given node) and sampling time. When constructing haplotype network, we follow the four main criteria: (1) mutation spectrum history. Mutations in ancestry haplotype should be contained in descendant haplotype; (2) minimum evolution (ME). The optimal haplotype network for a set of haplotypes is an arborescence whose sum of weights of all directed edges is minimal; (3) large ancestry haplotype. The probability of outward propagation is high for larger haplotype, and it is more likely to be an intermediate ancestor sequence node; (4) ancestry earlier sampling time. The sampling time of ancestral mutants is usually earlier than that of derived mutants.

The detailed construction process, (1) the directed edges representing the candidate immediate ancestor–descendant relationships were established according to the criterion that mutations in descendant should contain all mutations in its ancestor; (2) the immediate ancestor–descendant relationships were determined by the minimum evolution (ME) criterion in the field of phylogeny, which states that “the optimal phylogeny for a set of taxa is the one whose sum of edge weights is minimal” (Catanzaro, 2009), that is the sum of distances of all direct edges is minimized (the definition of distance between two haplotypes will be described later); (3) In practice, the haplotype networks may not be unique, so we proposed “large ancestry haplotypes and early ancestry haplotypes” criterion to determine which is the best among all haplotype networks.

The detailed algorithm:

First, let $h_{0},h_{1},...,h_{m-1}$ be all haplotypes, $m$ be the number of haplotypes and $V=\{h_{0},h_{1},...,h_{m-1}\}$ be the set of all haplotypes. Let $M=\{mut_{i}|mut_{i}=(pos_{i},ref_{i},alt_{i}),i\in I\}$ be the set of mutations including all mutation in haplotypes, where $pos_{i}\in\mathbb{Z}_{+}$ represents the genomic position of mutation$mut_{i}$, $ref_{i}$ is a reference nucleotide base or a sub-sequence of the reference sequence, $alt_{i}$ is the mutated nucleotide base. Let $M_{i}$ be subset of mutations set $M$, representing the mutation set of haplotypes $h_{i},i=0,1,\ldots,m-1$, respectively. According to the “mutation evolution criterion”, the set of candidates of directed edges should be

$E=\left\{ e_{ij}=\left( h_{i},h_{j} \right) | h_{i},h_{j}\in V,M_{i}\subset M_{j} \right\}.$	(1)

Second, the distance between any two haplotypes is defined as the number of sites of different mutations between the two haplotypes, e.g., the distance between haplotypes $h_{i}$ and $h_{j}$ is

$d_{ij}=\left| \left\{ pos | \exists ref,alt,s.t.\left( pos,ref,alt \right)\in\left( M_{i}\cup M_{j}-M_{i}\cap M_{j} \right) \right\} \right|$,	(2)

where $M_{i}$ and $M_{j}$ are sets of mutations of haplotypes $h_{i}$ and $h_{j}$, respectively. $\left| \cdot\right|$ denotes the cardinality of a set. Let $w\left( \cdot\right)$ be the cost function (or weight function) on $E$, $w:E\to\mathbb{Z}_{+}, e_{ij}\to d_{ij}$. By “Minimum evolution (ME) criterion”, the haplotype network must be the minimum-cost arborescence of weighted directed graph$G=\left( V,E,w \right)$. The difficulty of solving minimum-cost arborescence problem is determining the root of arborescence and handling directed cycles in graph. Fortunately, the root of minimum-cost arborescence should be the haplotype containing potential ancestor of the global pandemic and there is no directed cycle in $G$ because no mutation set of any haplotypes is strictly contained in itself. Suppose that $h_{0}$ is the haplotype containing potential haplotype ancestor. Let index set of directed edges candidates $E$ be $\Omega=\left\{ \left( i,j \right)|e_{ij}\in E \right\}$ and

$x_{ij}=\{\begin{aligned} 0,if\text{ }e_{ij}\notin arborescence \\ 1,if\text{ }e_{ij}\in arborescence \end{aligned}$.

The minimum-cost arborescence problem can be simplified as an integer programming problem:

$\begin{aligned} \min_{x_{ij}}\sum_{i,j} \tilde{d}_{ij}x_{ij} \\ \\ s.t.\sum_{i} x_{ij}\leq1 \\ \sum_{i,j} x_{ij}=m-1 \\ \tilde{d}_{ij}=\{\begin{aligned} d_{ij},M_{i}\subset M_{j} \\ +\infty,M_{i}\not\subset M_{j} \end{aligned} \\ x_{ij}\in\{0,1\} \end{aligned}$,	(3)

where the first constraint represents that the number of ancestors of any haplotype should be < 1, while the second constraint represents that the total number of edges in haplotype network must be m-1. $\tilde{d}_{ij}$ is set to $+\infty$ for $M_{i}\not\subset M_{j}$ according to the “mutation spectrum history criterion”.

Optimization problem (3) can be solved by keeping the closest ancestry of each haplotype except for the one containing the potential haplotype ancestor of the global pandemic. But there is a problem that the minimum-cost arborescence of $G$ may not be unique. If there are more than one haplotype which is both closest ancestry of a given haplotype, then the haplotype containing more sequences is chosen to be the ancestry of the given haplotype by “large ancestry haplotype criterion”. If there are more than one haplotype which is both closest ancestry of a given haplotype and is both meeting “large ancestry haplotype criterion”*,* then the one meeting the “ancestry earlier sampling time criterion” will be kept. Let $\prec$ be a relationship on 3-dimensional real space$\left\{ \left( x_{1},x_{2}{,x}_{3} \right) \right\}$, where $x_{1},x_{2}{,x}_{3}$ are real numbers. For any $a=\left( d_{1},{size}_{1},t_{1} \right), b=\left( d_{2},{size}_{2},t_{2} \right)$, $a\prec b$ if and only if $d_{1}<d_{2} \mathrm{or} \left( d_{1}=d_{2} \mathrm{and} {size}_{1}<{size}_{2} \right) \mathrm{or} (d_{1}=d_{2} \mathrm{and} {size}_{1}={size}_{2} \mathrm{and} t_{1}<t_{2})$. The overall problem would be

$\begin{aligned} \underset{x_{ij}}{\min_{\prec}}(\sum_{i,j} \tilde{d}_{ij}x_{ij},\sum_{i,j} size_{i}x_{ij},\sum_{i,j} \tilde{t}_{ij}x_{ij}) \\ s.t.\sum_{i} x_{ij}\leq1 \\ \sum_{i,j} x_{ij}=m-1 \\ \tilde{d}_{ij}=\{\begin{aligned} d_{ij},M_{i}\subset M_{j} \\ +\infty,M_{i}\not\subset M_{j} \end{aligned} \\ \tilde{t}_{ij}=\{\begin{aligned} 1,t_{i}\leq t_{j} \\ 0,t_{i}>t_{j} \end{aligned} \\ x_{ij}\in\{0,1\} \end{aligned}$,	(4)

where $t_{i}$ is the sampling time of $i$th haplotype $h_{i}$, ${size}_{i}$ represents the number of samples in haplotype $h_{i}$.

Let $t_{i},i=0,1,\ldots,m-1$ be the earliest sampling time of sequences in haplotype$h_{i},i=0,1,\ldots,m-1$, $size_{i},i=0,1,\ldots,m-1$ be the number of sequences in haplotype$h_{i},i=0,1,\ldots,m-1$. To solve problem (4), we first sort all haplotypes by the number of mutations and sequences (in descending order), and the earliest sampling time (in ascending order). Let $s$ be a permutation of set $\left\{ 0,1,\ldots,m-1 \right\}$, such that $\left| M_{s\left( i \right)} \right|>\left| M_{s\left( j \right)} \right| \mathrm{or} \left( \left| M_{s\left( i \right)} \right|=\left| M_{s\left( j \right)} \right| \mathrm{and} {size}_{s\left( i \right)}>{size}_{s\left( j \right)} \right) \mathrm{or} \left( \left| M_{s\left( i \right)} \right|=\left| M_{s\left( j \right)} \right| \mathrm{and} {size}_{s\left( i \right)}={size}_{s\left( j \right)} \mathrm{and} t_{s\left( i \right)}\leq t_{s\left( j \right)} \right),\mathrm{for} i<j.$ Then, find directed ancestor for each haplotype $h_{s\left( i \right)}$: for each $h_{s\left( j \right)}, j=i+1,i+2,\ldots,m-1$, find the smallest $j$ satisfying $M_{s\left( i \right)}\supset M_{s\left( j \right)}$. $h_{s\left( j \right)}$ would be the directed ancestor of $h_{s\left( i \right)}$. By sorting the haplotypes, McAN will reduce the time consumption of calculating distance between haplotypes. The time complexity of McAN is $O\left( m^{2} \right)$ which is equal to the time complexity of MSN and smaller than the time complexities of MJN ($O\left( m^{2.2} \right)$) (Bandelt, et al., 1999) and TCS ($O\left( m^{5} \right)$) (Clement, et al., 2002), where $m$is the number of haplotypes.

**Text. S2. Generation of simulated dataset.**

The simulated dataset were generated using JC69 model (Jukes and Cantor, 1969). Specifically, the common ancestor sequence with 1000 nucleobase is generated randomly, and two descendants for each sequence were generated until the 10th generation via JC69 model by setting the parameter $\alpha$ = 0.0001 which represents the probability of change from one nucleotide to a different one.

**Figures**

**Fig. S1. Illustration of four criterions adopted by McAN for haplotype network construction.**

**Fig. S2. Overview of McAN algorithm.** (A) Haplotypes were sorted by the number of mutations in descending order first, the number of sequences in descending order second, and the earliest sampling time in ascending order third. Mutation 1A means nucleobase in position 1 is replaced by A; (B) Illustration of constructing haplotype network via McAN; (C) McAN finds the ancestor of each haplotype directly instead of calculating distance matrix.

**Fig. S3. Haplotype networks of SARS-CoV-2 sequences from Diamond Princess (DP) dataset via MSN, MJN, TCS and McAN.** 482 SARS-CoV-2 sequences including 70 sequences collected from diamond princess cruise ship and 412 sequences collected globally available on GISAID as of March 10, 2020 were used for constructing haplotype network. First, haplotype network was constructed via McAN. Then, MSN, MJN and TCS implemented in PopART were used to construct haplotype network as comparison.

**Fig. S4. Haplotype network of SARS-CoV-2 lineages constructed by McAN.** 130 major haplotypes of sublineages of L and S lineages from 121,618 SARS-CoV-2 genomes were used for constructing haplotype network via McAN.

**Fig. S5. Haplotype networks of monkeypox and human influenza A viruses constructed by McAN.** (A) Haplotype network of 44 monkeypox sequences constructed by McAN; (B) Haplotype network of 21 human influenza A viruses (IAV) sequences of the hemagglutinin (HA) gene from 1918 flu strains constructed by McAN.

**Fig. S6. The number of haplotypes in terms of the number of sequences.** The number of haplotypes in terms of the number of SARS-CoV-2 sequences ranging from a hundred thousand to one million.
