## Supplementary Figures for "McAN: an ultrafast haplotype network construction algorithm"

### Figure S1

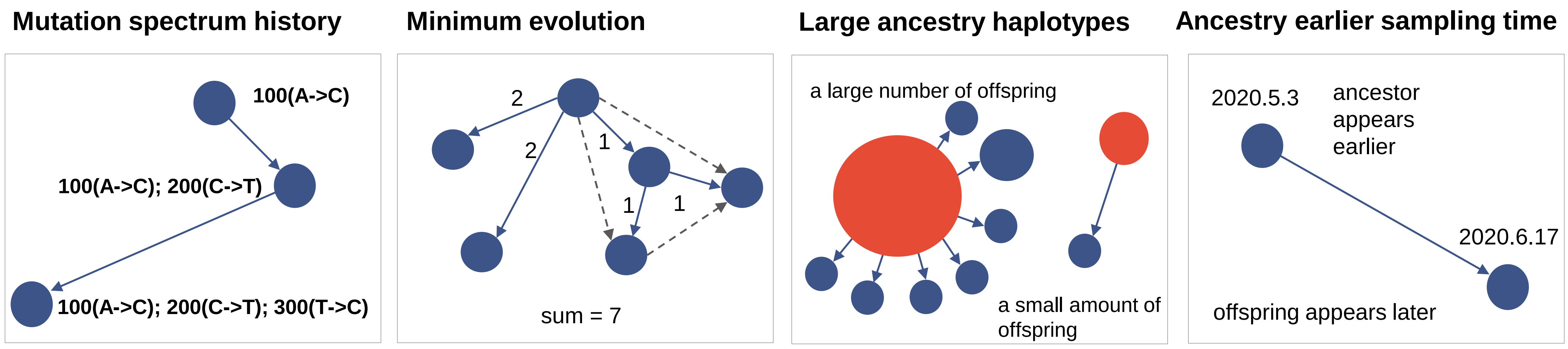

Mutation spectrum history
100(A->C)
100(A->C); 200(C->T)
100(A->C); 200(C->T); 300(T->C)
Minimum evolution
2
2
1
1
1
sum = 7
Large ancestry haplotypes
a large number of offspring
a small amount of offspring
Ancestry earlier sampling time
2020.5.3
ancestor appears earlier
2020.6.17
offspring appears later

### Figure S2

A

| Haplotype | Mutations | Number of Samples | Earliest sampling time |
|---|---|---|---|
| Hap_0 | (1A,3C) | 3 | 2020.07.04 |
| Hap_1 | (1A,2G) | 1 | 2020.02.21 |
| Hap_2 | (2G) | 2 | 2020.02.05 |
| Hap_3 | (1A) | 2 | 2020.05.19 |
| Hap_4 (root) | () | 10 | 2020.01.02 |

B

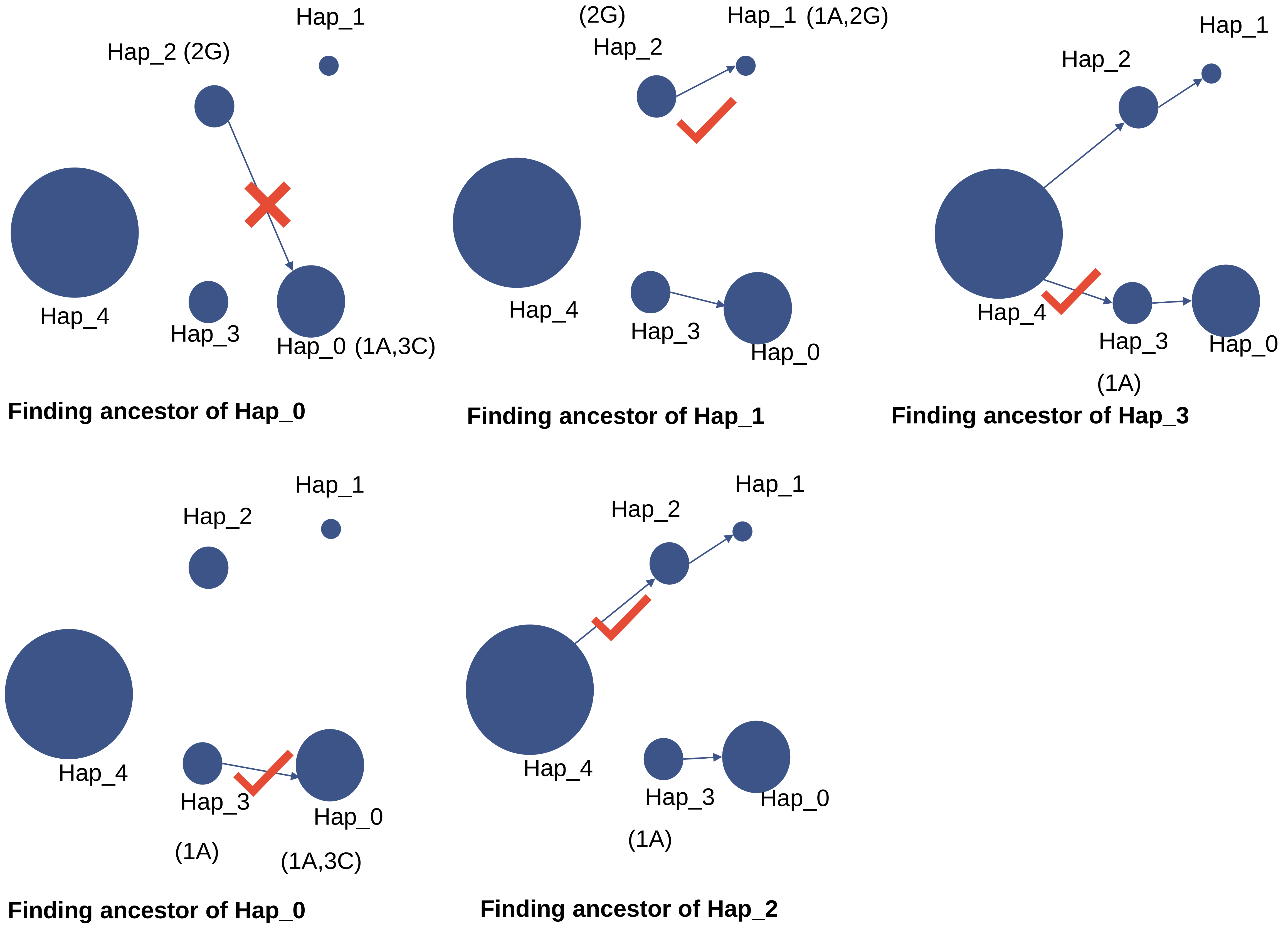


C

| | Hap_1 (1A,2G) | Hap_2 (2G) | Hap_3 (1A) | Hap_4 () |
|---|---|---|---|---|
| Hap_0 (1A,3C) | | X | V | |
| Hap_1 (1A,2G) | | V | | |
| Hap_2 (2G) | | | | V |
| Hap_3 (1A) | | | | V |

**McAN**
Finding the ancestors for only **5** times and only restore the **4** ancestors in memory

| | Hap_1 (1A,2G) | Hap_2 (2G) | Hap_3 (1A) | Hap_4 () |
|---|---|---|---|---|
| Hap_0 (1A,3C) | 2 | 3 | 1 | 2 |
| Hap_1 (1A,2G) | | 1 | 1 | 2 |
| Hap_2 (2G) | | | 2 | 1 |
| Hap_3 (1A) | | | | 1 |

**Existing algorithms**
Calculate distance between haplotypes for **10** times and restore all the **10** distance in memory

#### Figure S3

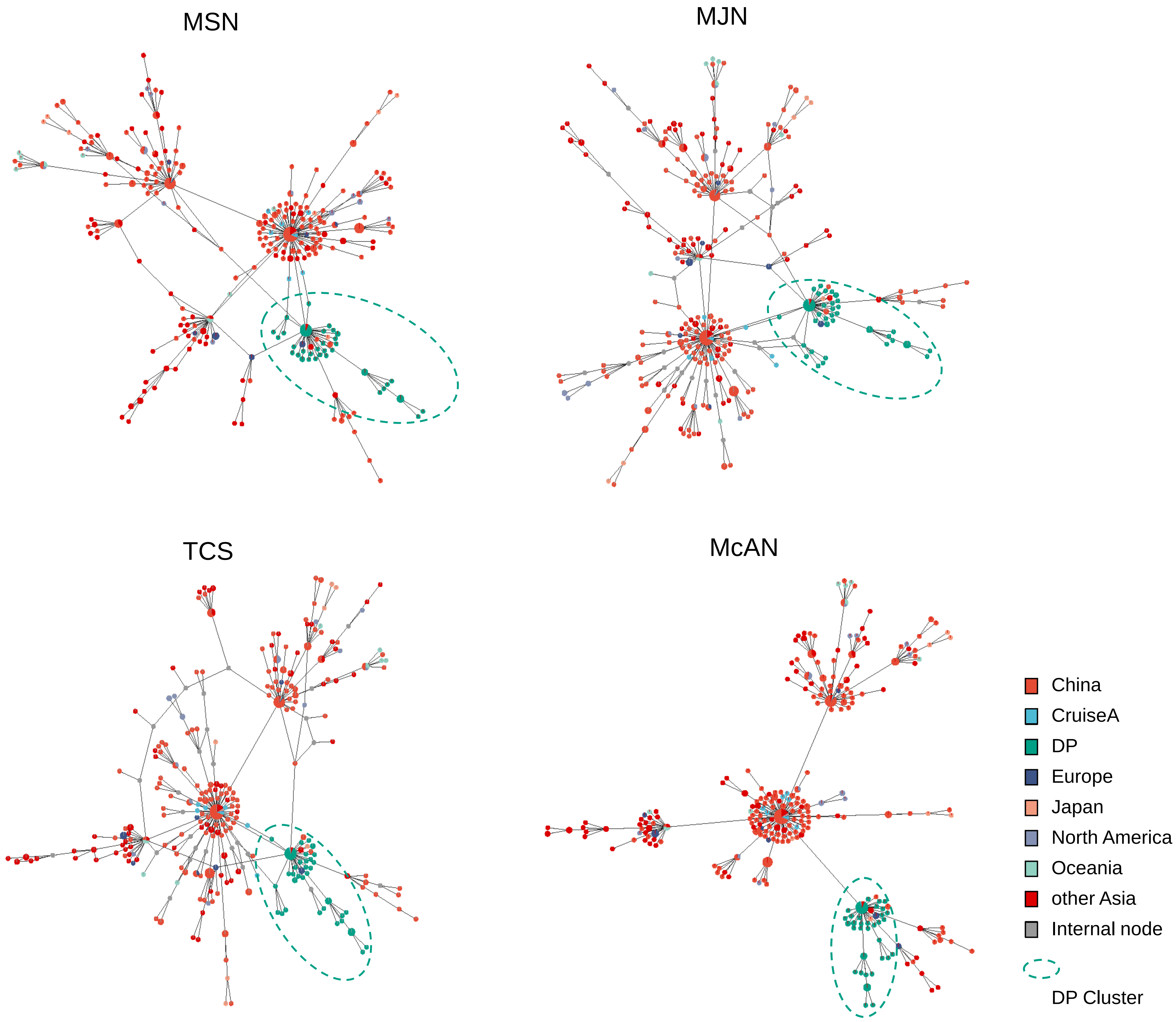

MSN
MJN
TCS
McAN
China
CruiseA
DP
Europe
Japan
North America
Oceania
other Asia
Internal node
DP Cluster

### Figure S4

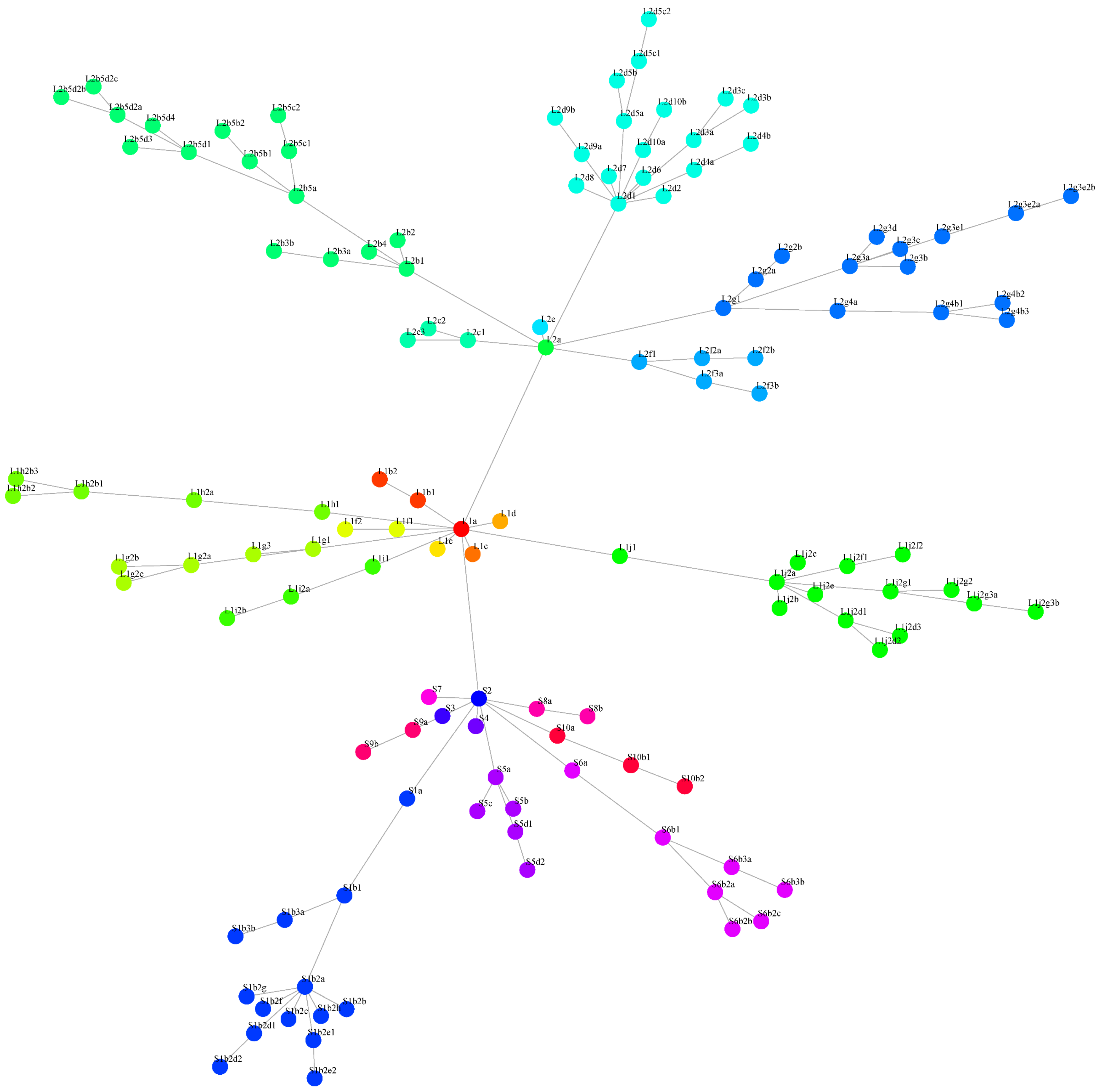
L2d5c2
L2d5c1
L2d5b
L2d5a
L2d9b
L2d9a
L2d8
L2d7
L2d6
L2d1
L2d2
L2d10b
L2d10a
L2d3c
L2d3b
L2d3a
L2d4b
L2d4a
L2b5d2b
L2b5d2c
L2b5d2a
L2b5d4
L2b5d3
L2b5d1
L2b5b2
L2b5b1
L2b5c2
L2b5c1
L2b5a
L2b4
L2b2
L2b3b
L2b3a
L2b1
L2c2
L2c3
L2c1
L2e
L2a
L2f1
L2f2a
L2f2b
L2f3a
L2f3b
L2g1
L2g2a
L2g2b
L2g3a
L2g3d
L2g3c
L2g3b
L2g3e1
L2g3e2a
L2g3e2b
L2g4a
L2g4b1
L2g4b2
L2g4b3
L1h2b3
L1h2b2
L1h2b1
L1h2a
L1h1
L1b2
L1b1
L1f2
L1f1
L1a
L1d
L1e
L1c
L1g1
L1g3
L1g2a
L1g2b
L1g2c
L1i1
L1i2a
L1i2b
L1j1
L1j2a
L1j2b
L1j2c
L1j2e
L1j2f1
L1j2f2
L1j2g1
L1j2g2
L1j2g3a
L1j2g3b
L1j2d1
L1j2d2
L1j2d3
S7
S2
S3
S4
S8a
S8b
S9a
S9b
S10a
S10b1
S10b2
S5a
S5b
S5c
S5d1
S5d2
S6a
S6b1
S6b2a
S6b2b
S6b2c
S6b3a
S6b3b
S1a
S1b1
S1b3a
S1b3b
S1b2a
S1b2g
S1b2f
S1b2c
S1b2h
S1b2b
S1b2d1
S1b2d2
S1b2e1
S1b2e2

### Figure S5

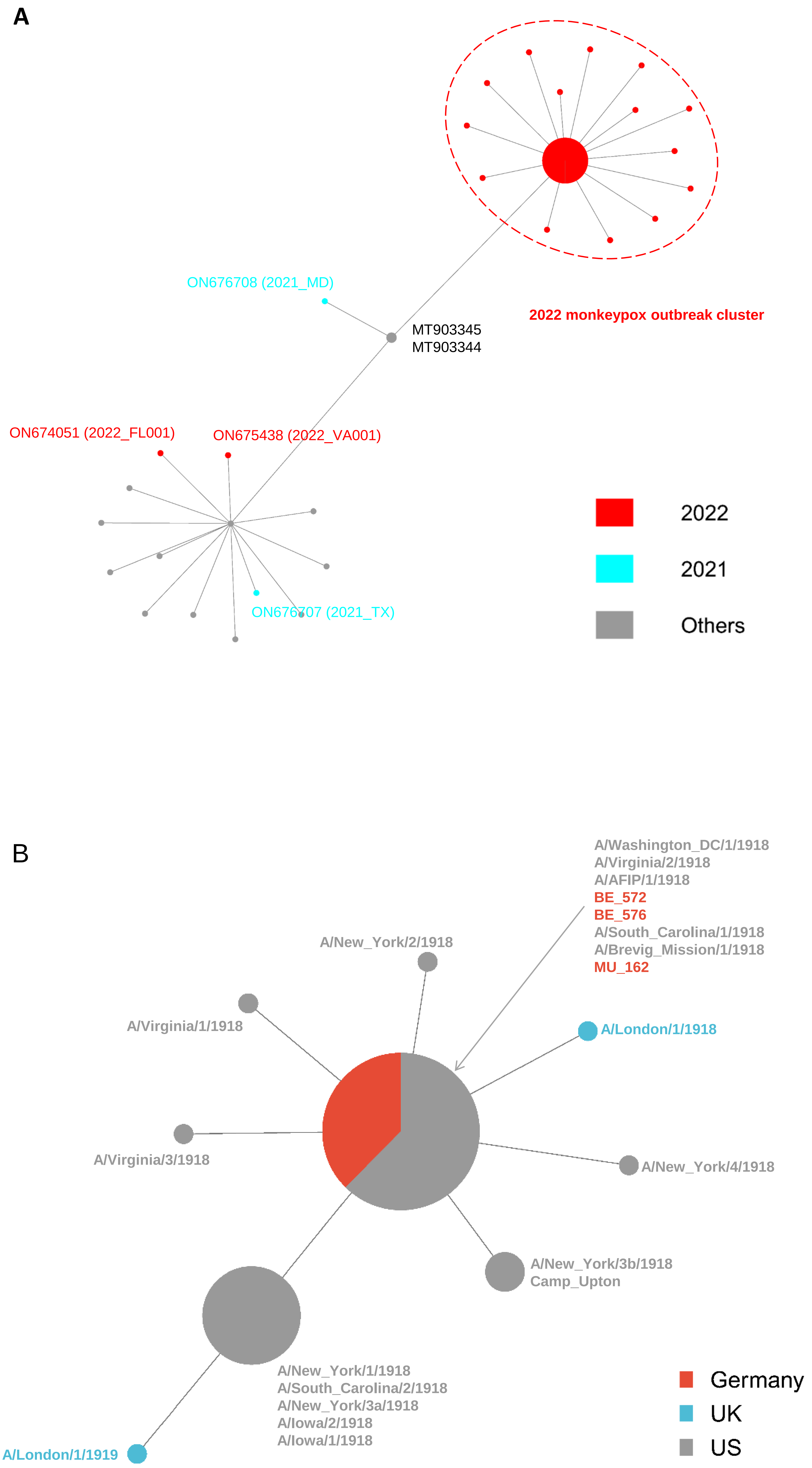
A
ON676708 (2021_MD)
MT903345
MT903344
2022 monkeypox outbreak cluster
ON674051 (2022_FL001)
ON675438 (2022_VA001)
ON676707 (2021_TX)
2022
2021
Others
B
A/Washington_DC/1/1918
A/Virginia/2/1918
A/AFIP/1/1918
BE_572
BE_576
A/South_Carolina/1/1918
A/Brevig_Mission/1/1918
MU_162
A/New_York/2/1918
A/Virginia/1/1918
A/London/1/1918
A/Virginia/3/1918
A/New_York/4/1918
A/New_York/3b/1918
Camp_Upton
A/New_York/1/1918
A/South_Carolina/2/1918
A/New_York/3a/1918
A/Iowa/2/1918
A/Iowa/1/1918
A/London/1/1919
Germany
UK
US

**Figure S6**

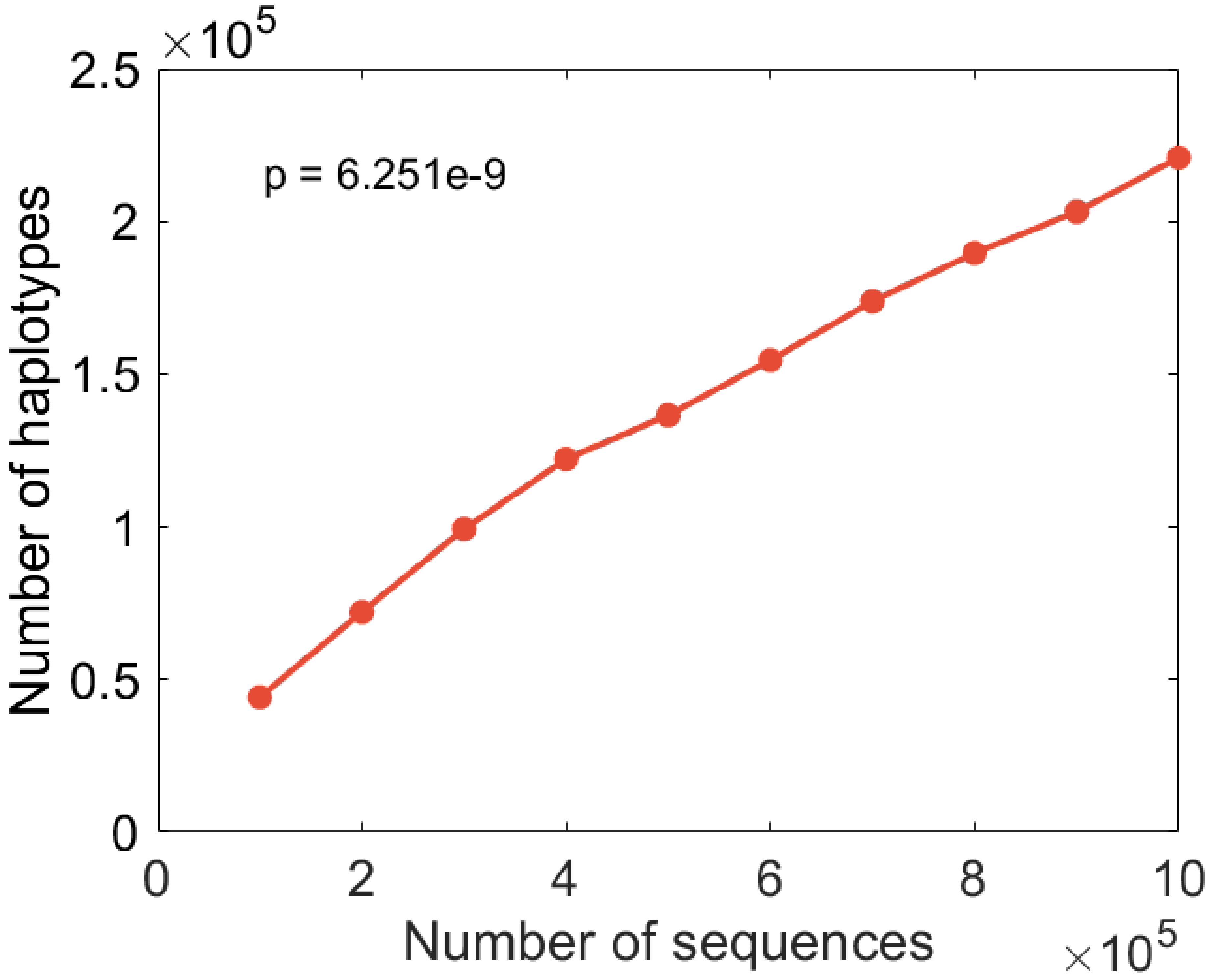
×10^5
2.5
2
1.5
1
0.5
0
p = 6.251e-9
Number of haplotypes
0
2
4
6
8
10
Number of sequences
×10^5
